# Hybridization led to a rewired pluripotency network in the allotetraploid *Xenopus laevis*

**DOI:** 10.1101/2022.09.14.507817

**Authors:** Wesley A. Phelps, Matthew D. Hurton, Taylor N. Ayers, Anne E. Carlson, Joel C. Rosenbaum, Miler T. Lee

## Abstract

After fertilization, maternally contributed factors to the egg initiate the transition to pluripotency to give rise to embryonic stem cells, in large part by activating de novo transcription from the embryonic genome. Diverse mechanisms coordinate this transition across animals, suggesting that pervasive regulatory remodeling has shaped the earliest stages of development. Here, we show that maternal homologs of mammalian pluripotency reprogramming factors OCT4 and SOX2 divergently activate the two subgenomes of *Xenopus laevis*, an allotetraploid that arose from hybridization of two diploid species ~18 million years ago. Although most genes have been retained as two homeologous copies, we find that a majority of them undergo asymmetric activation in the early embryo. Chromatin accessibility profiling and CUT&RUN for modified histones and transcription factor binding reveal extensive differences in enhancer architecture between the subgenomes, which likely arose through genomic disruptions as a consequence of allotetraploidy. However, comparison with diploid *X. tropicalis* and zebrafish shows broad conservation of embryonic gene expression levels when divergent homeolog contributions are combined, implying strong selection to maintain dosage in the core vertebrate pluripotency transcriptional program, amid genomic instability following hybridization.

## Introduction

In mammals, zygotic genome activation (ZGA) is triggered after an initial period of transcriptional quiescence, during the slow first cleavages post fertilization (Svoboda, 2018). This is a few days removed from the subsequent induction of pluripotent stem cells in the blastocyst by a core network of factors including NANOG, OCT4 and SOX2 (Li and Belmonte, 2017; Takahashi and Yamanaka, 2016). In contrast, faster-dividing taxa including zebrafish, *Xenopus*, and Drosophila activate their genomes in the blastula hours after fertilization during the maternal-to-zygotic transition (MZT) (Foe and Alberts, 1983; Kane and Kimmel, 1993; Newport and Kirschner, 1982a; Vastenhouw et al., 2019), which leads immediately to pluripotency. In zebrafish, maternally provided homologs of NANOG, OCT4 and SOX2 are required for genome activation (Lee et al., 2013; Leichsenring et al., 2013; Miao et al., 2022); thus, vertebrate embryos deploy conserved pluripotency induction mechanisms at different times during early development.

Beyond vertebrates, unrelated maternal factors direct genome activation and the induction of stem cells, e.g. Zelda (Liang et al., 2008), CLAMP (Colonnetta et al., 2021; Duan et al., 2021) and GAF (Gaskill et al., 2021) in Drosophila, though they seem to share many functional aspects with vertebrate pluripotency factors, including pioneering roles in opening repressed embryonic chromatin and establishing activating histone modifications (Blythe and Wieschaus, 2016; Gaskill et al., 2021; Hug et al., 2017). This diversity of strategies implies that the gene network regulating pluripotency has been extensively modified over evolutionary time (Endo et al., 2020; Fernandez-Tresguerres et al., 2010), though it is unknown when and under what circumstances major modifications arose.

We sought to understand how recent genome upheaval has affected the pluripotency regulatory network in the allotetraploid *Xenopus laevis*, by deciphering how embryonic genome activation is coordinated between its two subgenomes. *X. laevis’s* L (long) and S (short) subgenomes are inherited from each of two distinct species separated by ~34 million years that hybridized ~18 million years ago (Session et al., 2016) (Fig. 1A). A subsequent whole-genome duplication restored meiotic pairing. Despite extensive rearrangements and deletions, most genes are still encoded as two copies (homeologs) on parallel, non-inter-recombining chromosomes (Session et al., 2016). Previously, homeologs had been challenging to distinguish due to high functional and sequence similarity; however, the recent high-quality *X. laevis* genome assembly has made it feasible to resolve differential expression and regulation genome-wide between the two subgenomes (Elurbe et al., 2017; Session et al., 2016).

**Fig. 1.**
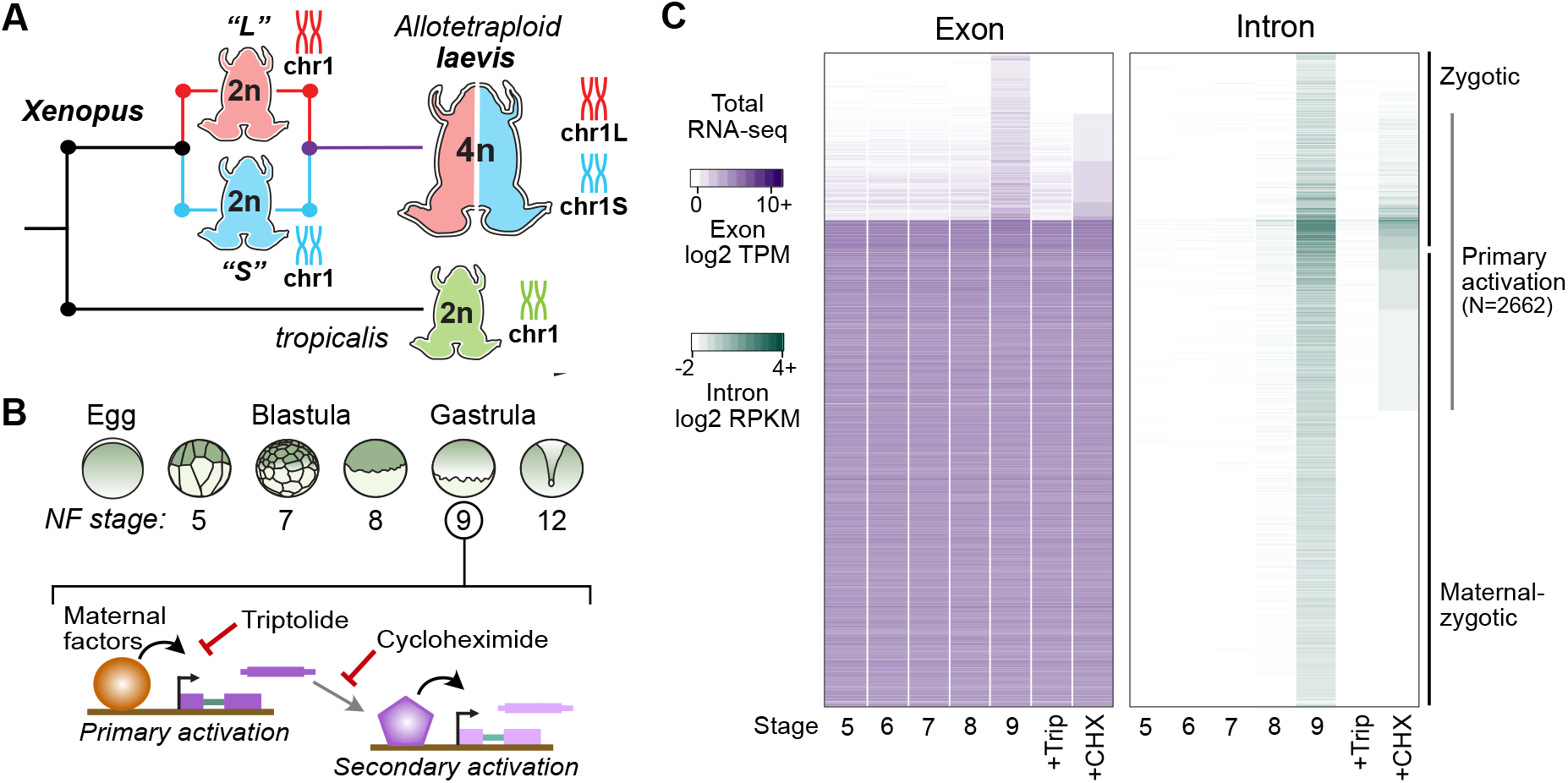
Identifying the first wave of genome activation across the two subgenomes. (**A**) The allotetraploid *X. laevis* genome contains two distinct subgenomes “L” and “S” due to interspecific hybridization of ancestral diploids. (**B**) Triptolide inhibits genome activation, as measured in the late blastula, while cycloheximide inhibits only secondary activation, distinguishing genes directly activated by maternal factors. NF = Nieuwkoop and Faber. (**C**) Heatmap of RNA-seq coverage over exons (left) and introns (right) of activated genes.

Allopolyploidy often provokes acute effects on gene expression (Hu and Wendel, 2019; Moran et al., 2021), leading to regulatory shifts over time to reconcile dosage imbalances and incompatibilities between gene copies (C. E. Grover et al., 2012; Song et al., 2020; Swamy et al., 2021). This phenomenon has been explored primarily in plants (Adams and Wendel, 2005; Husband et al., 2013; Mable, 2004), but the extent to which this has occurred in the few characterized allopolyploid vertebrates is unclear (Chen et al., 2019b; Li et al., 2021; Luo Jing et al.). For *X. laevis*, there is a broad trend toward balanced homeolog expression across development and adult tissues (Session et al., 2016) and an overall ontogenetic and transcriptomic trajectory similar to 48-million-years diverged diploid *X. tropicalis* (Harland and Grainger, 2011; Yanai et al., 2011). However, initial observations suggest a divergent cis-regulatory landscape between the two *X. laevis* subgenomes (Elurbe et al., 2017; Ochi et al., 2017).

Although *Xenopus* embryos have long been a model for understanding the MZT, e.g. (Amodeo et al., 2015; Charney et al., 2017; Chen et al., 2019a; Gentsch et al., 2019; Gibeaux et al., 2018; Gurdon et al., 1958; Kimelman et al., 1987; Newport and Kirschner, 1982b, 1982a; Paraiso et al., 2019; Skirkanich et al., 2011; Veenstra et al., 1999; Yanai et al., 2011), ZGA regulators have not previously been identified in *X. laevis*. Here, we elucidate the top-level regulators of *X. laevis* pluripotency and ZGA, and the enhancer architecture that differentially recruits them to homeologous gene copies between the two subgenomes. Despite differential subgenome activation, combined transcriptional output converges to proportionally resemble the diploid state, maintaining gene dosage for the embryonic pluripotency program.

### Identifying divergently activated homeologous genes

At genome activation, the *X. laevis* pluripotency network consists of maternal regulators acting directly on the first embryonic genes (Fig. 1B). To identify these genes, we performed a total RNA-seq early embryonic time course using our *X. laevis*-specific ribosomal RNA depletion protocol (Phelps et al., 2021) (Fig. 1A,B, Supplementary Table 1). We identified 4772 genes with significant activation by the middle of Nieuwkoop and Faber (N.F.) stage 9 (8 hours post fertilization [h.p.f.] at 23°C) (Fig. 1C, Supplementary Table 2), through a combination of exon- and intron-overlapping sequencing reads deriving from nascent pre-mRNA (Lee et al., 2013). Indeed, two-thirds of these genes had substantial maternal contributions that masked their activation when quantifying exon-overlapping reads alone (Fig. 1C). These genes fail to be activated in embryos treated at 1-cell stage with the transcription inhibitor triptolide (Gibeaux et al., 2018) when compared to DMSO vehicle control embryos (Fig. 1B,C, Supplementary Fig. 1A-C).

To distinguish direct targets of maternal factors (primary activation) (Fig. 1B), we then performed RNA-seq on stage 9 embryos treated with cycloheximide at stage 8, to inhibit translation of newly synthesized embryonic transcription factors that could regulate secondary activation (Harvey et al., 2013; Lee et al., 2013). 2662 genes (56% of all activated genes) were still significantly activated in cycloheximide-treated embryos compared to triptolide-treated embryos, representing the first wave of genome activation in the embryo (Fig. 1C, Supplementary Fig. 1A).

We analyzed subgenome of origin for activated genes and found that they are preferentially encoded as two homeologous copies in the genome (*P* = 2.9×10^-181^, χ-squared test, 6 d.o.f.) (Fig. 2A). However, a majority of these genes have asymmetric expression between the two homeologs, often with transcription deriving from only the L or S copy alone (Fig. 2B-C, Supplementary Fig. 1D). This degree of divergent activation suggests large differences in the cis-regulatory architecture between gene homeologs in the two subgenomes. Genes activated from both subgenomes are enriched in transcriptional regulators (*P* < 0.01, Fisher’s exact test, two-sided) (Supplementary Fig. 1E), suggesting that gene function may have influenced homeolog expression patterns. However, there is no evidence for strong functional divergence between homeologs expressed asymmetrically between the subgenomes, as estimated by non-synonymous versus synonymous mutation rate in coding regions (dN/dS ratio) (Supplementary Fig. 1F,G).

**Fig. 2.**
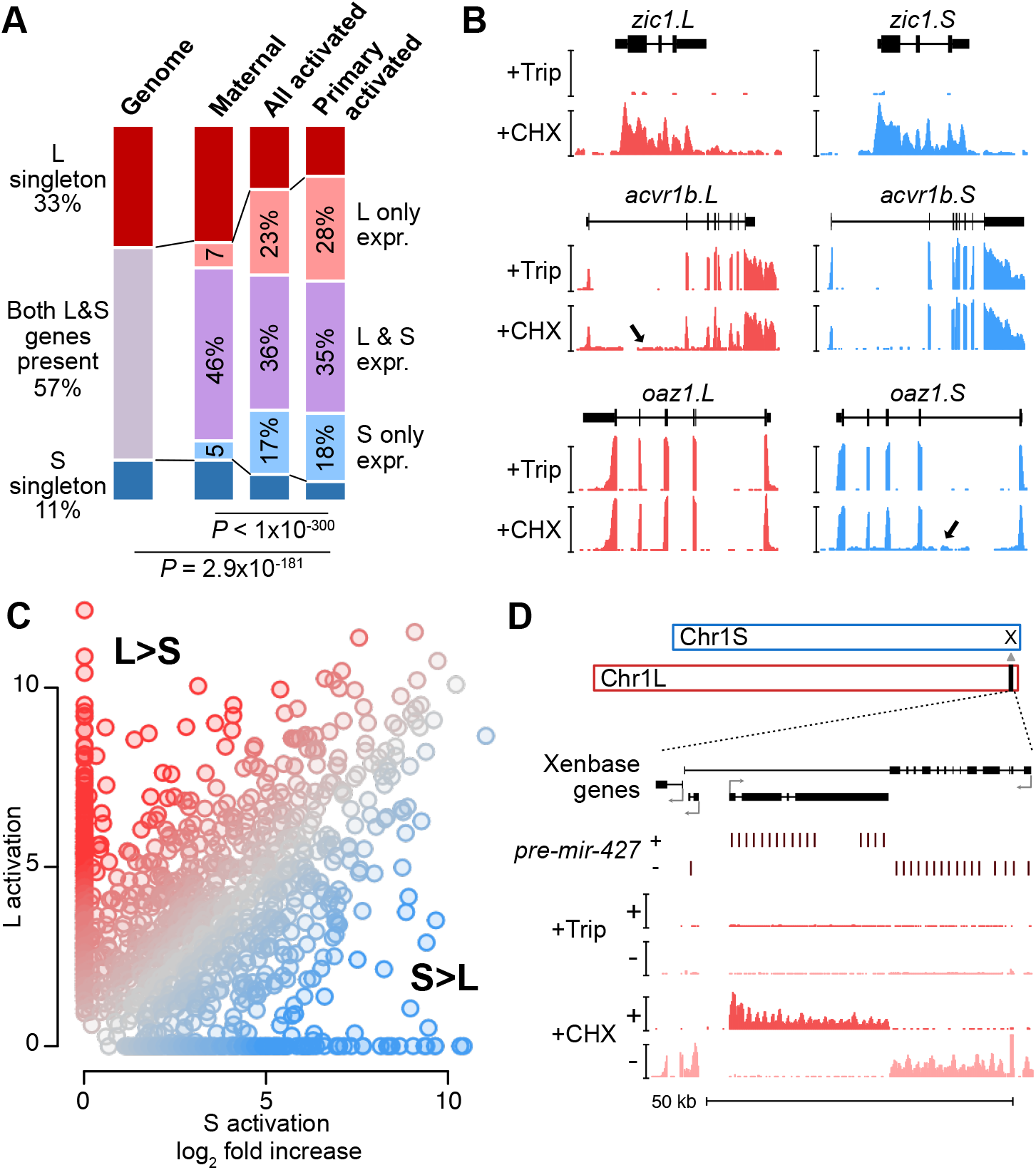
Homeologous genes are differentially activated in the early embryo. (**A**) Proportion of genes encoded as homeologs on both subgenomes versus only one subgenome (singleton) (left), as compared to expression patterns in the early embryo. *P* = 2.9×10^-181^, χ-squared test, 6 d.o.f., comparing genomic to expressed proportions; *P* < 1×10^-300^, χ-squared test, 8 d.o.f., comparing proportions within expressed genes. (**B**) Browser tracks showing log2 reads-per-million RNA-seq coverage of equivalently activated homeologs (top) and differentially activated homeologs (L-specific, middle; S-specific, bottom). (**C**) Biplot comparing log2 fold activation of homeologs in cycloheximide versus triptolide treated embryos. (**D**) Browser track showing strand-separated reads-per-million RNA-seq coverage over the *mir-427* encoding locus on the distal end of Chr1L (v10.1). Trip = triptolide, CHX = cycloheximide.

### The microRNA *mir-427* is encoded on only one subgenome

Among the first-wave genes is the microRNA *mir-427*, which plays a major role in clearance of maternally contributed mRNA (Lund et al., 2009). Similar to *X. tropicalis mir-427* (Owens et al., 2016) and the related zebrafish *mir-430* (Lee et al., 2013), *mir-427* is one of the most strongly activated genes in the *X. laevis* embryonic genome (Fig. 2D, Supplementary Fig. 1C, 2A). In version 9.2 of the *X. laevis* genome assembly, the *miR-427* precursor hairpin sequence is found in only five copies overlapping a Xenbase-annotated long non-coding RNA on chr1L (Supplementary Fig. 2B). This is in stark contrast to the 171 tandemly arrayed precursors in the two *X. tropicalis mir-427* loci on Chr03, which is thought to accelerate mature *miR-427* accumulation during the MZT to facilitate rapid maternal clearance (Owens et al., 2016). Zebrafish similarly encodes a large array of 55 *mir-430* precursors, which begin to target maternal mRNA for clearance shortly after ZGA (Bazzini et al., 2012; Giraldez et al., 2006; Lee et al., 2013).

To better capture the genomic configuration of the *mir-427* primary transcript, we aligned the miRBase-annotated precursor sequence (Kozomara et al., 2019) to the recently released version 10.1 *X. laevis* genome assembly. This revealed an expanded *mir-427* locus at the distal end of Chr1L composed of 33 precursor copies, encoded in both strand orientations over 55 kilobases (Fig 2D, Supplementary Fig. 2A). This is reminiscent of the *X. tropicalis* configuration (Owens et al., 2016), though smaller in scale and on a non-homologous chromosome. The corresponding region on Chr1S is unalignable (Supplementary Fig. 2C), suggesting that *mir-427* is encoded on only the L subgenome. We additionally found two *mir-427* hairpin sequence matches to the distal end of Chr3S, but these loci were not supported by substantial RNA-seq coverage (Supplementary Fig. 2D). These results strongly suggest that the mir-427 locus has undergone genomic remodeling, resulting in absence from the S subgenome, but possibly also translocation between chromosomes between the *tropicalis* and *laevis* lineages.

### Subgenomes differ in their regulatory architecture

To discover the maternal regulators of differential homeolog activation, we first profiled embryonic chromatin using Cleavage Under Target & Release Using Nuclease (CUT&RUN) (Hainer and Fazzio, 2019; Skene and Henikoff, 2017), which we adapted for blastulae. We found that cell dissociation was necessary for efficient nuclear isolation to carry out the on-bead CUT&RUN chemistry (Fig. 3A, Supplementary Fig. 3A-C). At stages 8 and 9, the active marks H3 lysine 4 trimethylation (H3K4me3) and H3 lysine 27 acetylation (H3K27ac) were enriched in activated genes compared to their unactivated homeologs (stage 8 H3K27ac, P < 3×10^-8^; stage 8 H3K4me3, *P* < 0.01; stage 9 H3K4me3, *P* < 2×10^-10^, paired t-tests, two-sided) (Fig. 3B,C, Supplementary Fig. 3D, Supplementary Table 3). Differential promoter engagement by transcriptional machinery likely underlies the differential active histone levels; however, we found no promoter sequence differences between homeologs that would implicate differential recruitment of specific transcription factors (Supplementary Table 4).

**Fig. 3.**
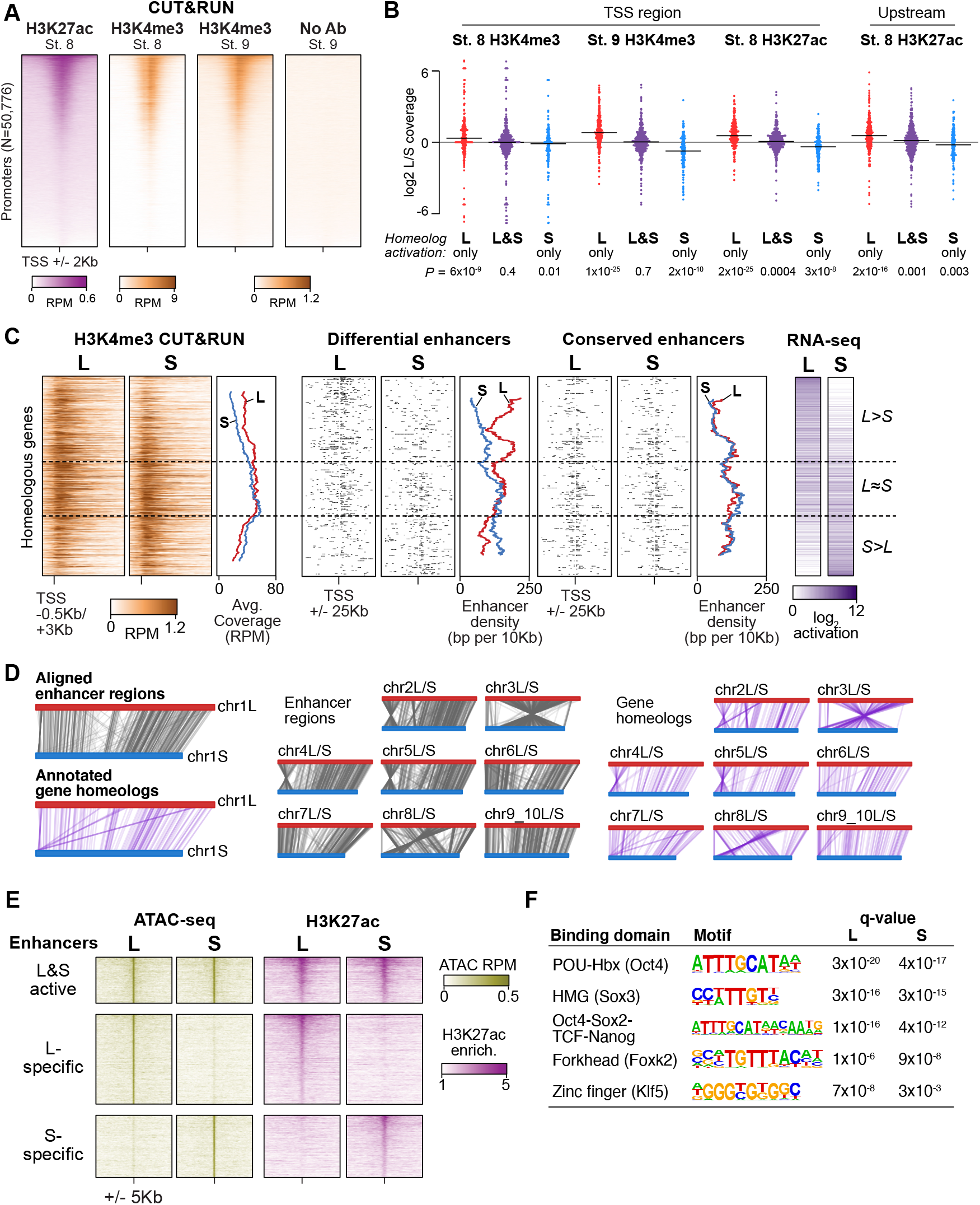
Differential homeolog activation is regulated by subgenome-specific enhancers. (**A**) CUT&RUN coverage over all annotated transcription-start site (TSS) regions, sorted by descending stage 8 H3K27ac signal. (**B**) Bee-swarm plots showing the log2 ratio of L versus S homeolog coverage among genes where only one homeolog is activated (L only, S only), or both homeologs are activated. TSS region is 1 kb centered on the TSS; upstream region is 500 bp to 3 kb upstream of the TSS. Horizontal bars show medians. *P* values are from two-sided paired t-tests of log2 L homeolog coverage vs log2 S homeolog coverage. (**C**) Stage 9 H3K4me3 CUT&RUN coverage over paired homeologous gene regions around the TSS (left) and maps comparing enhancer density near homeologous TSSs (middle). Differential enhancers are active in only one subgenome, conserved enhancers are active in both. Average densities are plotted to the right of each paired map. Gene pairs are sorted according to L versus S subgenome RNA-seq activation ratio (right). (**D**) Schematics showing aligned enhancers and their homeologous regions (gray) mapped onto L (red, top lines) and S (blue, bottom lines) chromosomes. Comparable schematics show Xenbase annotated homeologous gene pairs (lavender). (**E**) Heatmap of stage 9 ATAC-seq and stage 8 H3K27ac CUT&RUN over L & S homeologous regions for equivalently active enhancers (top) and subgenome-specific enhancers. (**F**) Top enriched transcription factor motif families in L-specific and S-specific active enhancers compared to inactive homeologous regions. FDR-corrected *P*-values from Homer are shown. RPM = reads per million.

Instead, we searched for differences in gene-distal regulatory elements – i.e., enhancers – between the two subgenomes. To identify regions of open chromatin characteristic of enhancers, we performed Assays for Transposase-Accessible Chromatin with sequencing (ATAC-seq) on dissected animal cap explants from stage 8 and 9 embryos; the high concentration of yolk in vegetal cells inhibits the Tn5 transposase (Esmaeili et al., 2020). We called peaks of elevated sub-nucleosome sized fragment coverage, then intersected the open regions with our H3K27ac CUT&RUN. This yielded 7562 putative open and acetylated enhancers at genome activation (Supplementary Fig. 3E, Supplementary Table 5).

To identify homeologous L and S enhancer regions, we constructed a subgenome chromosome-chromosome alignment using LASTZ (Harris, 2007). This yielded a syntenic structure consistent with genetic maps (Fig. 3D) (Session et al., 2016), recapitulating the large inversions between chr3L/chr3S and chr8L/chr8S. 79% of enhancer regions successfully lifted over to homeologous chromosomes, and of these, >90% of these are flanked by the same homeologous genes (Supplementary Fig. 3F), confirming local synteny.

Among the paired regions, only 23% had conserved enhancer activity in both homeologs, with the remaining pairs exhibiting differential H3K27ac and chromatin accessibility (Fig. 3E, Supplementary Fig. 3G). Differential enhancer density around genes significantly correlated with differential activation (*P* = 1.3×10^-16^, Pearson’s correlation test) (Fig. 3C, middle), with greater L enhancer density around differentially activated L genes, and similarly for S enhancers and S genes. In contrast, conserved enhancers had equivalent density near both homeologs regardless of activation status (*P* = 0.20, Pearson’s correlation test) (Fig. 3C, right). Thus, differences in enhancer activity likely underlie divergent gene homeolog transcription at genome activation.

### Maternal pluripotency factors differentially engage the subgenomes

Given that these paired enhancer regions are differentially active despite having similar base sequences, we searched for transcription factor binding motifs that distinguished active enhancers from their inactive homeolog. Two motifs were strongly enriched in both active L enhancers and active S enhancers, corresponding to the binding sequences of the pluripotency factors OCT4 and SOX2/3 (SOXB1 family) (Fig. 3F, Supplementary Table 4). Since mammalian OCT4 and SOX2 are master regulators of pluripotent stem cell induction (Takahashi and Yamanaka, 2016), and zebrafish homologs of these factors are maternally provided and required for embryonic genome activation (Lee et al., 2013; Leichsenring et al., 2013; Miao et al., 2022), we hypothesized that differential enhancer binding by maternal *X. laevis* OCT4 and SOXB1 homologs underlies asymmetric activation of the L and S subgenomes.

RNA-seq reveals high maternal levels of *pou5f3.3* (OCT4 homolog) and *sox3* mRNA, each deriving from both subgenomes (Supplementary Fig. 3H). To assess their roles in genome activation, we inhibited their translation using previously validated antisense morpholinos (Morrison and Brickman, 2006; Zhang et al., 2003) injected into stage 1 embryos. Each of the two morpholinos was complementary to both the L and S homeologs of *pou5f3.3* and *sox3*, respectively, but not to their paralogs that are primarily expressed zygotically (i.e., *pou5f3.1* and weakly maternal *pou5f3.2*). To again focus specifically on maternal regulation of primary genome activation, we treated the injected embryos with cycloheximide at stage 8 and collected them at stage 9 for RNA-seq. When both Pou5f3.3 and Sox3 were inhibited, we observed significant downregulation of 62% of activated genes compared to embryos injected with a control morpholino, including the *mir-427* transcript (Fig. 4A, Supplementary Fig. 4A). Targeting *pou5f3.3* or *sox3* mRNA individually had minimal impact on genome activation (Supplementary Fig. 4A), suggesting these two maternal transcription factors together coordinate early gene expression.

**Fig. 4.**
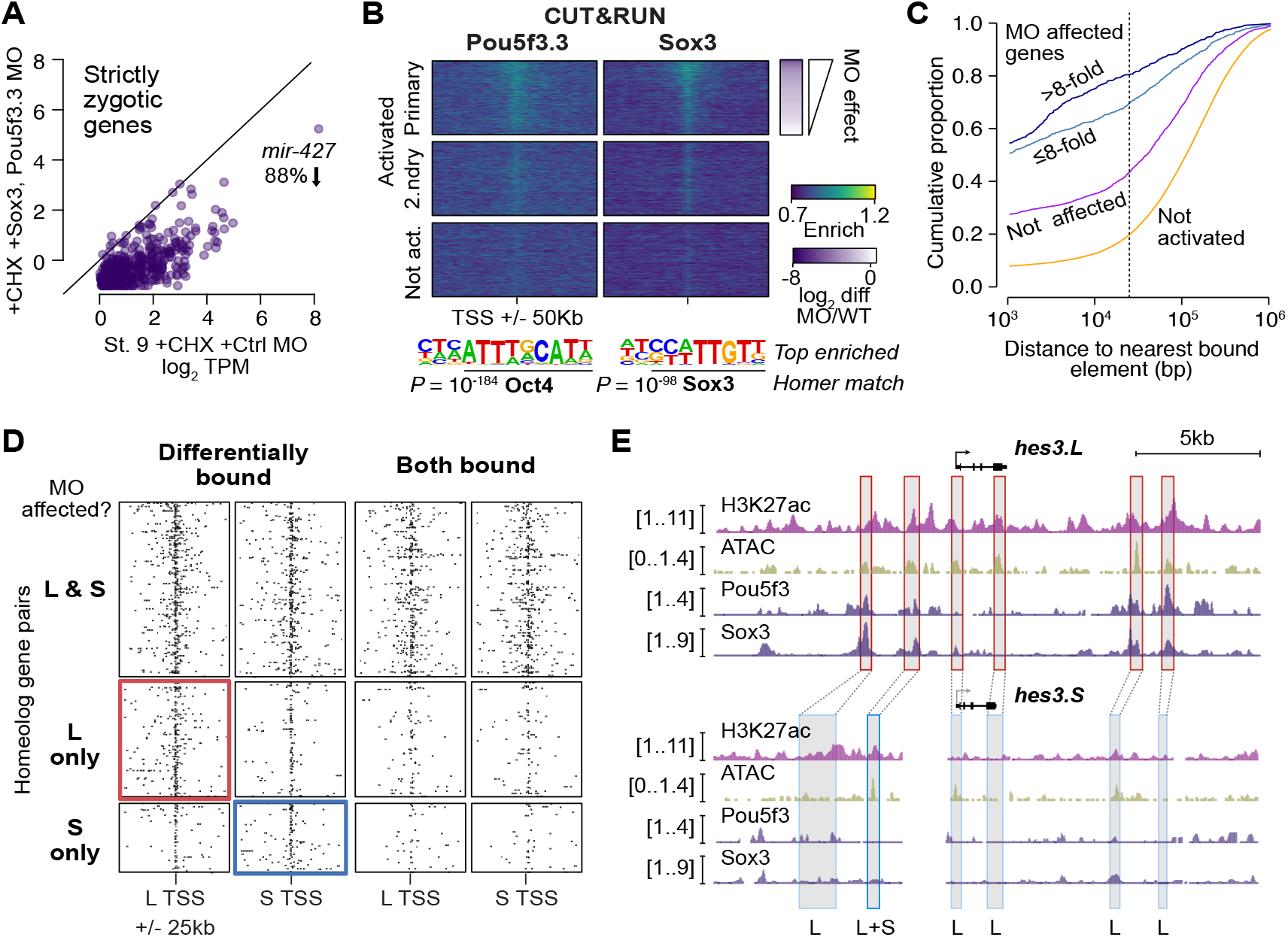
Pou5f3.3 and Sox3 binding drives genome activation. (**A**) Biplot showing inhibited gene activation in *pou5f3.3/sox3* morpholino-treated embryos compared to controls. (**B**) Stage 8 Pou5f3.3 (left) and Sox3 (right) CUT&RUN coverage near TSSs for genes activated in the primary and secondary waves (top, middle) and unactivated genes (bottom). Primary activated genes are sorted by RNA-seq sensitivity to *pou5f3.3/sox3* morpholino. Top enriched motifs for each factor are shown below. (**C**) Cumulative distributions of distance from a Pou5f3/Sox3-bound regulatory element for genes strongly (>8-fold) and less strongly affected by *pou5f3.3/sox3* morpholino compared to unaffected and unactivated genes. (**D**) Maps showing density of Pou5f3/Sox3-bound regulatory elements around paired homeologous TSSs, divided into elements with differential homeologous L & S binding (left panels) versus both bound (right panels). TSSs are grouped according to L versus S homeolog sensitivity to *pou5f3.3/sox3* morpholino treatment. (**E**) Browser tracks showing CUT&RUN enrichment and ATAC-seq coverage near active homeolog *hes3.L* and inactive homeolog *hes3.S*. One shared enhancer (L+S) and five L-specific regulatory regions are highlighted.

To interrogate Pou5f3.3 and Sox3 chromatin binding across the subgenomes, we performed CUT&RUN on stage 8 embryos injected at stage 1 with mRNA encoding V5 epitope-tagged *pou5f3.3.L* and *sox3.S*. Peak calling revealed thousands of binding sites for each factor (Supplementary Fig. 4B-F), and Homer de novo motif analysis recovered the OCT4 and SOX3 binding sequences as top hits (*P* = 10^-184^ and *P* = 10^-98^, respectively) (Fig. 4B, Supplementary Fig. 4G,H). CUT&RUN signal for both factors is enriched in the vicinity of activated genes, with stronger association to genes affected by morpholino treatment (*P* < 1×10^-300^, Kruskal-Wallis test) (Fig. 4B,C, Supplementary Fig. 4I), and indeed comparison between differentially affected homeolog pairs showed preferential binding in enhancers near the Pou5f3.3/Sox3-dependent homeolog (Pou5f3.3: *P* = 1.5×10^-5^; Sox3: *P* = 1.8×10^-5^, Kruskal-Wallis tests) (Fig. 4D,E, Supplementary Fig. 4J,K). Together, these results implicate Pou5f3.3 and Sox3 in regulating ZGA differentially between the two subgenomes.

### The ancestral pluripotency program is maintained, despite enhancer turnover

Finally, to understand differential activation given the natural history of *X. laevis* allotetraploidy, we compared *X. laevis* subgenome activation patterns to diploid *X. tropicalis* as a proxy for the ancestral *Xenopus*, since there are no known extant diploid descendants of either *X. laevis* progenitor (Session et al., 2016). For three-way homeologs/orthologs with minimal maternal contribution in *X. laevis*, there is broad conservation of relative expression levels between the *X. tropicalis* and *X. laevis* embryonic transcriptomes after genome activation, when *X. laevis* homeolog levels are summed gene-wise (Pearson’s *r* = 0.72) (Fig. 5A, left, Supplementary Table 6). However, the correlation weakens when the *X. laevis* subgenomes are considered independently: relative activation levels in one subgenome alone are depressed relative to *X. tropicalis*, with expression of some genes completely restricted to one subgenome or the other (L, Pearson’s *r* = 0.60; S, Pearson’s *r* = 0.55) (Fig. 5A, middle, right). If the diploid L and S progenitor embryos each exhibited the inferred ancestral activation levels, then these trends strongly suggest that *X. laevis* underwent regulatory remodeling post allotetraploidization that maintained relative gene expression dosage for embryonic genome activation.

**Fig. 5.**
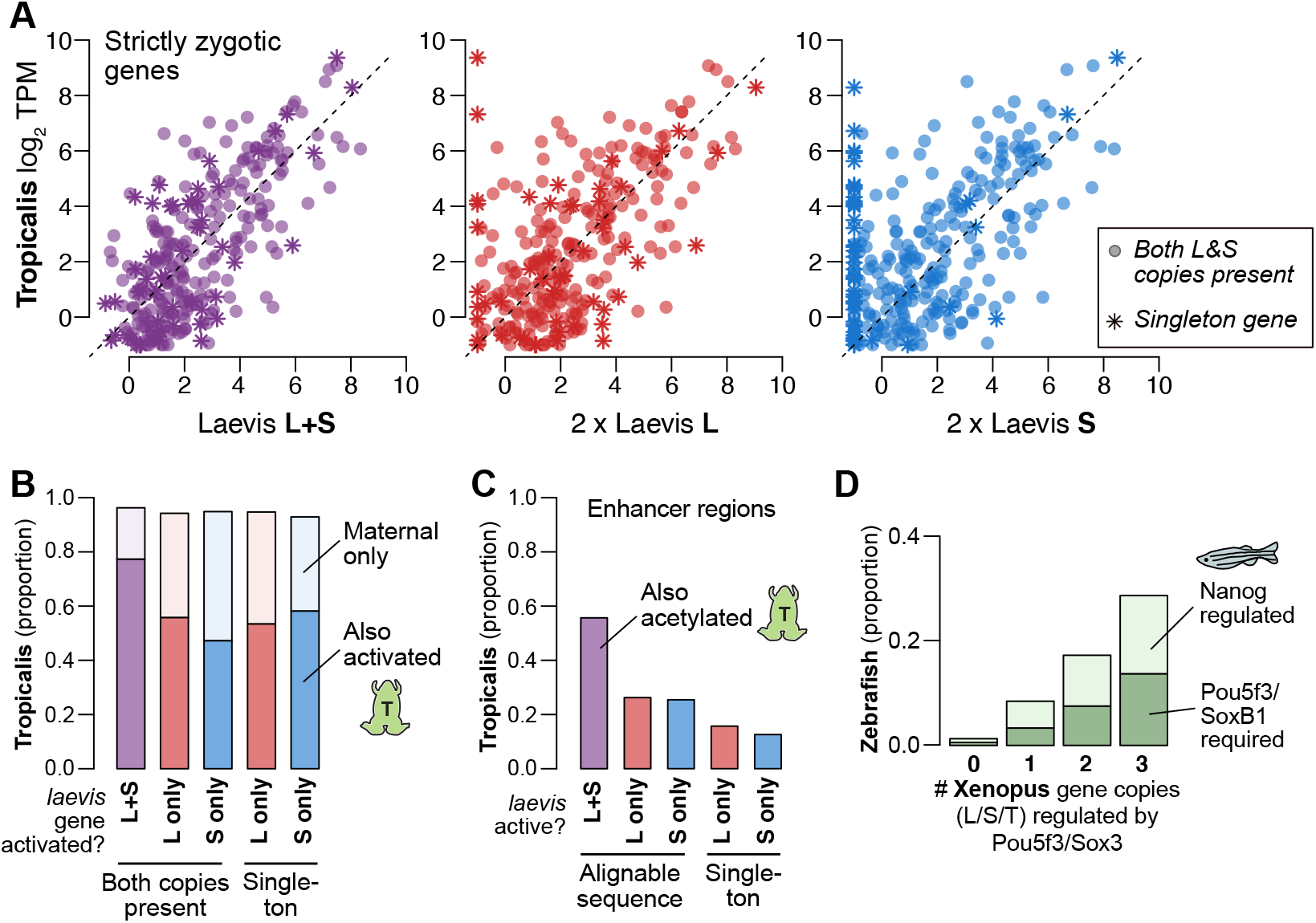
Regulatory divergence underlies dosage maintenance. (**A**) Biplots comparing relative expression levels of activated genes in *X. laevis* and *X. tropicalis*, treating L and S homeolog contributions separately (middle, right) or summed (left). Individual subgenome expression is scaled 2x, since transcript per million (TPM) normalization is calculated relative to the entire *X. laevis* transcriptome. (**B**) Barplots showing the proportion of *X. laevis* genes across activation categories whose orthologs are also activated in *X. tropicalis* or part of the maternal contribution. (**C**) Barplots showing the proportion of *X. laevis* enhancers across activity categories that are acetylated in *X. tropicalis*. (**D**) Barplots showing the proportion of *Xenopus* genes whose orthologs are regulated by Pou5f3/SoxB1 and Nanog in zebrafish. *Xenopus* genes are classified according to how many homeo/orthologs are regulated by Pou5f3/Sox3. Genes with conserved regulation in both *X. laevis* homeologs and *X. tropicalis* are more likely to be regulated by Pou5f3/SoxB1 in zebrafish, but also more likely to be regulated by Nanog.

However, most differentially activated genes also have a maternal contribution, which could offset asymmetries in homeolog activation levels. Indeed, overall when both *X. laevis* gene homeologs are activated, the *X. tropicalis* ortholog is more likely also to be activated, compared to genes where only one homeolog is activated (*P* = 5.0×10^-20^, χ-squared test, 4 d.o.f.) (Fig. 5B), suggesting a greater degree of regulatory innovation among differentially activated homeologs. Indeed, enhancers conserved between the *X. laevis* subgenomes exhibit significantly higher conservation with *X. tropicalis*, versus subgenome-specific enhancers (*P* = 1.0×10^-300^, χ-squared test, 4 d.o.f.) (Fig. 5C, Supplementary Fig. 5A). However, total embryonic expression (i.e., maternal + zygotic) appears to be broadly maintained between *X. laevis* and *X. tropicalis* (Fig. 5B), suggesting that much of the divergent subgenome activation is buffered by the maternal contribution, maintaining the stoichiometry of mRNA in the embryonic transcriptome.

This trend is also apparent at greater evolutionary distances. We find that genes activated in *X. laevis* are largely also expressed in zebrafish embryos (~450 million years separated) (Supplementary Fig. 5B). Despite considerable divergence in activation timing, coactivated *X. laevis* homeologs are still more likely to be part of the first wave of zebrafish genome activation (*P* = 8.0×10^-12^, χ-squared test, 4 d.o.f.) and targeted by maternal homologs of OCT4 and SOX2, but also NANOG (*P* = 1.5×10’^138^, χ-squared test, 6 d.o.f.) (Fig. 5D, Supplementary Fig. 5C-E). Interestingly, *Xenopus* and possibly all Anuran amphibians lack a NANOG ortholog, likely due to a chromosomal deletion (Schuff et al., 2012). In the absence of a Nanog homolog in the maternal contribution, we find that maternal Pou5f3.3 and Sox3 seem to have subsumed NANOG’s roles in *X. laevis* genome activation, while zygotic factors such as Ventx help promote cell potency in the early gastrula (Scerbo et al., 2012; Schuff et al., 2012). This demonstrates core-vertebrate mechanistic conservation in genome activation amid both cis- and trans-regulatory shuffling, which converge to support pluripotent stem cell induction and embryonic development.

## Discussion

Together, our findings establish the pluripotency factors Pou5f3.3 and Sox3 as maternal activators of embryonic genome activation, which are differentially recruited to the two homeologous subgenomes of *X. laevis* by a rewired enhancer network (Fig. 6). Of the thousands of genes activated during the MZT, a majority of annotated homeolog pairs experience differential activation, which appears to be driven by subgenome-specific enhancer gain and/or loss correlated with differential Pou5f3.3/Sox3 binding and regulation. However, this magnitude of regulatory divergence seems to have had a net neutral effect, as combined subgenome activation produces a composite reprogrammed embryonic transcriptome akin to diploid *X. tropicalis*.

**Fig. 6.**
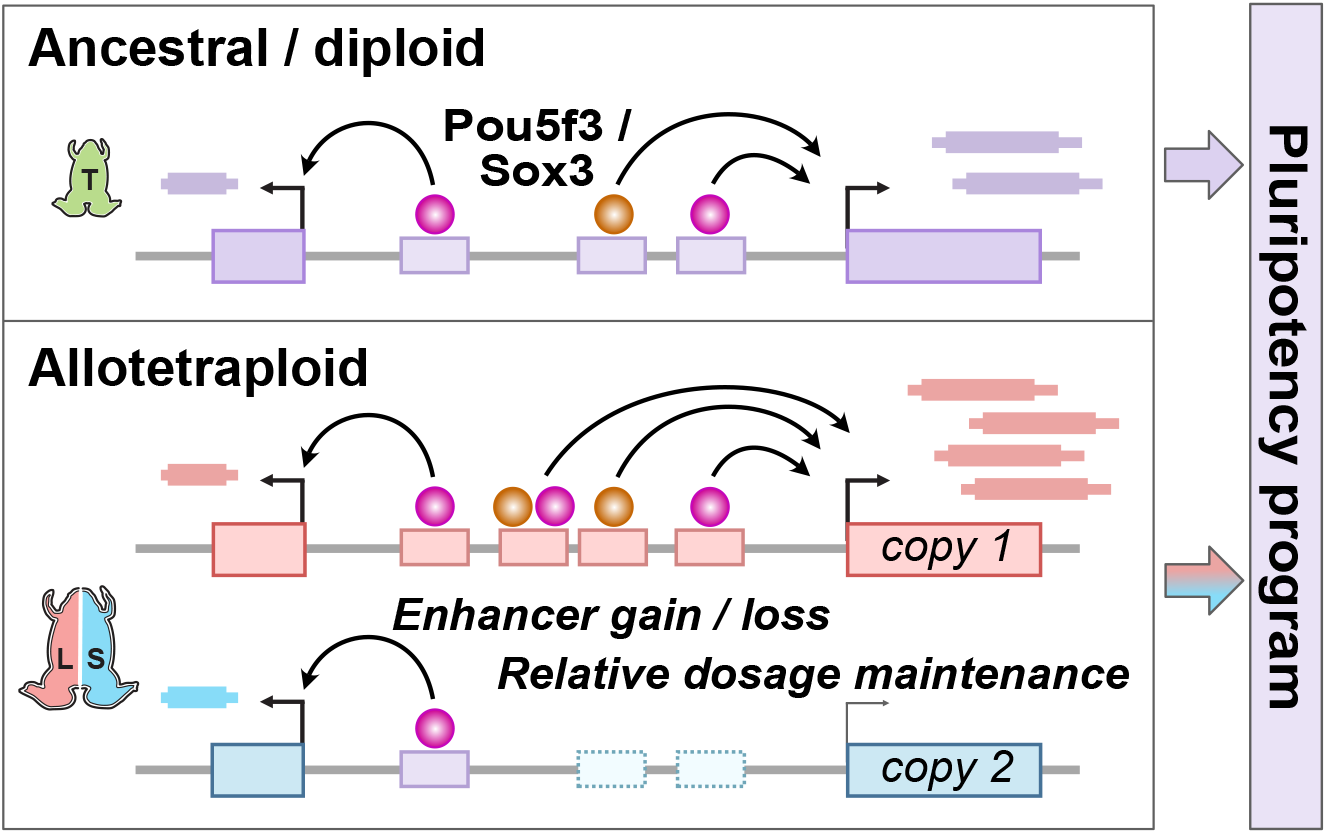
Model for pluripotency network evolution. *X. laevis* likely underwent extensive enhancer turnover between its two subgenomes, which nonetheless maintained stoichiometry of pluripotency reprogramming in the early embryo.

As embryogenesis proceeds, regulatory divergence between the subgenomes is likely even broader. In *X. tropicalis*, signal transducers and transcription factors including Pou5f3.2/3, Sox3, Smad1/2, β-catenin, Vegt, Otx1, and Foxh1 regulate embryo-wide and regional gene activation (Charney et al., 2017; Gentsch et al., 2019; Paraiso et al., 2019), and binding motifs for some of these are found in differentially active *X. laevis* enhancers (Fig. 3F, Supplementary Table 4). Additionally, by focusing on accessible chromatin in animal caps, we may have underestimated the magnitude of homeologous enhancer divergence regulating endodermal fate in the vegetal cells. But based on the close morphological similarity of *X. tropicalis* and *X. laevis* embryos, we would predict that these subgenome regulatory differences also converge to producing ancestral dosages in the transcriptome.

Although homeolog expression bias can derive from gene regulatory differences evolved in the parental species prior to hybridization (Buggs et al., 2014; C. E. Grover et al., 2012), we propose that regulatory upheaval in *X. laevis* post-hybridization (i.e., “genome shock” (McClintock Barbara, 1984)) led to expression level gain or loss in one homeolog, which was subsequently corrected by compensatory changes to the other homeolog, possibly repeatedly (Shi et al., 2012; Tirosh Itay et al., 2009). This implies that early development exerts constraint on the reprogrammed embryonic transcriptome while tolerating (or facilitating) regulatory turnover. The apparent reconfiguration of the *mir-427* cluster after the *X. laevis* and *tropicalis* lineages split similarly highlights how essential MZT regulatory mechanisms can evolve, ostensibly neutrally given that *miR-427-directed* maternal clearance is conserved in *Xenopus*. Thus, *X. laevis* embryos illustrate how the pluripotency program may have accommodated regulatory network disruptions, genomic instability, and aneuploidy across the animal tree.

## Supplementary information

Supplementary Figures 1-5

Supplementary Table 1: RNA-seq expression values and differential expression analysis Supplementary Table 2: Activated homeologous genes

Supplementary Table 3: Promoter annotations

Supplementary Table 4: Sequence motif analyses

Supplementary Table 5: Enhancer annotations

Supplementary Table 6: Cross-species comparisons

## Data availability

Sequencing data are available in the Gene Expression Omnibus (GEO) under accession number GSE207027. Code and auxiliary data files are available on Github, github.com/MTLeeLab/xl-zga. Additional data files including chromosome alignments are available at OSF, osf.io/ct6g8/

## Author contributions

WAP & MTL conceived of the project, performed analyses, and wrote the manuscript. WAP performed the experiments with assistance from MDH, TNA, and JCR. AEC oversaw *Xenopus* egg collection.

## Acknowledgements

We thank S. Hainer and lab for providing the pAG-MNase enzyme, assistance with the CUT&RUN protocol, and feedback. We thank M. Rebeiz, T. Levin, M. Turcotte, K. Arndt and lab, C. Kaplan and lab, and the entire Lee lab for feedback. This project used the University of Pittsburgh Health Sciences Core at UPMC Children’s Hospital Pittsburgh for sequencing. This work was supported by the March of Dimes #5-FY16-307, the National Institutes of Health R35GM137973, the Samuel and Emma Winters Foundation, and start-up funds from the University of Pittsburgh to MTL, and by the University of Pittsburgh Center for Research Computing through the resources provided.

## Methods

### Animal Husbandry

All animal procedures were conducted under the supervision and approval of the Institutional Animal Care and Use Committee at the University of Pittsburgh. *Xenopus laevis* adults (Research Resource Identifier NXR_0.0031; NASCO) were housed in a recirculating aquatic system (Aquaneering) at 18°C with a 12/12 h light/dark cycle. Frogs were fed 3x weekly with Frog Brittle (NASCO #SA05960 (LM)M).

### Embryo Collection

Sexually mature females were injected with 1000 IU human chorionic gonadotropin into their dorsal lymph sac and incubated overnight at 16°C. Females were moved to room temperature to lay. Eggs from two mothers per collection were artificially inseminated using dissected testes in MR/3 (33 mM NaCl, 0.6 mM KCl, 0.67 mM CaCl_2_, 0.33 mM MgCl_2_, 1.67 mM HEPES, pH 7.8) (Hazel L Sive, 2000). Dissected testes were stored up to one week in L-15 medium at 4°C prior to use. Zygotes were de-jellied (Hazel L Sive, 2000) in MR/3 pH 8.5, with 0.3%β-mercaptoethanol with gentle manual agitation, neutralized with MR/3 pH 6.5, washed twice with MR/3 and incubated in MR/3 at 23°C until desired developmental stage based on morphology.

### RNA-seq libraries

All stage 9 embryos were collected halfway through the stage, at 8 hours post fertilization. Triptolide samples were bathed in 20 μM triptolide in DMSO (200X stock added to MR/3) at stage 1 and cycloheximide samples were bathed in 500 μg/mL cycloheximide in DMSO at the beginning of stage 8; both were collected when batch-matched, untreated embryos were halfway through stage 9. Equivalent volumes of DMSO were used to treat control samples. Previously validated morpholinos targeting *pou5f3.3* (GTACAATATGGGCTGGTCCATCTCC) (Morrison and Brickman, 2006) and *sox3* (AACATGCTATACATTTGGAGCTTCA) (Zhang et al., 2003) along with control GFP morpholino (ACAGCTCCTCGCCCTTGCTCACCAT) were ordered from GeneTools. Morpholino treated embryos were injected at stage 1 with *pou5f3.3, sox3*, and/or GFP control morpholino: 40 ng *pou5f3.3* + 40 ng GFP, 40 ng *sox3* + 40 ng GFP, 40 ng *pou5f3.3* + 40 ng sox3, or 80 ng GFP. High concentration morpholino injections were 55 ng *pou5f3.3* + 75 ng *sox3*. Each embryo was injected twice with 5 nl of MO on opposite sides. Embryos were allowed to recover to stage 5 before moving to MR/3 to develop, and collected when batch-matched, untreated embryos were halfway through stage 9.

For RNA extraction, two embryos per sample were snap frozen and homogenized in 500 μl of TRIzol Reagent (Invitrogen #15596026) followed by 100 μl of chloroform. Tubes were spun at 18,000 x g at 4°C for 15 minutes, the aqueous phase was transferred to a fresh tube with 340 μl of isopropanol and 1 μl of GlycoBlue (Invitrogen #AM9515), then precipitated at −20°C overnight. Precipitated RNA was washed with cold 75% ethanol and resuspended in 50 μl of nuclease-free water. Concentration was determined by NanoDrop.

For library construction, rRNA depletion was performed as per Phelps et al 2021 with *X. laevis* specific oligos reported previously (Phelps et al., 2021): 1 μl of antisense nuclear rRNA oligos and 1 μl of antisense mitochondrial rRNA oligos (final concentration 0.1 μM per oligo) were combined with 1 μg of total RNA in a 10 μl buffered reaction volume (100 mM Tris-HCl pH 7.4, 200 mM NaCl, 10 mM DTT), heated at 95°C for 2 minutes and cooled to 22°C at a rate of 0.1°C/s in a thermocycler. Next, 10U of thermostable RNaseH (NEB #M0523S) and 2 μl of provided 10X RNaseH buffer were added and volume brought to 20 μl with nuclease-free water. The reaction was incubated at 65°C for 5 or 30 minutes, then 5U of TURBO DNase (Invitrogen #AM2238) and 5 μl of provided 10x buffer was added, volume brought to 50 μl with nuclease-free water and incubated at 37C for 30 minutes. The reaction was purified and size selected to >200 nts using Zymo RNA Clean and Concentrator-5 (Zymo #R1013) according to manufacturer’s protocol, eluting in 10 μl of nuclease-free water. The WT Stage 5 sample was also depleted of mitochondrial COX2 and COX3 mRNA as part of the Phelps et al 2021 study. Strand-specific RNA-seq libraries were constructed using NEB Ultra II RNA-seq library kit (NEB #E7765) according to manufacturer’s protocol with fragmentation in first-strand buffer at 94°C for 15 minutes. Following first and second strand synthesis, DNA was purified with 1.8X AmpureXP beads (Beckman #A63880), end repaired, then ligated to sequencing adaptors diluted 1:5. Ligated DNA was purified with 0.9X AmpureXP beads and PCR amplified for 8 cycles, then purified again with 0.9X AmpureXP beads. Libraries were verified by Qubit dsDNA high sensitivity (Invitrogen #Q32851) and Fragment Analyzer prior to multiplexed sequencing at the Health Sciences Sequencing Core at Children’s Hospital of Pittsburgh.

### CUT&RUN

CUT&RUN procedure was adapted from Hainer et al (Hainer and Fazzio, 2019) optimizations of the method of Skene and Henikoff (Skene and Henikoff, 2017). For nuclear extraction, embryos were de-vitellinized using 1 mg/mL pronase dissolved in MR/3. Once the vitelline envelope was removed, 12 – 24 embryos (50K – 100K cells) were carefully transferred into 1 mL of NP2.0 buffer (Briggs et al., 2018) in a 1.5 mL tube and gently agitated (pipetting buffer over the surface of the embryos) until cells have dissociated. The buffer was carefully drawn off to the level of the cells and 1 mL of Nuclear Extraction (NE) buffer (20mM HEPES-KOH, pH 7.9, 10mM KCl, 500μM spermidine, 0.1% Triton X-100, 20% glycerol) with gentle pipetting with a clipped P1000, and the lysate was centrifuged at 600xg in 4°C for 3 min. The free nuclei were then bound to 300 μL of activated concanavalin A beads (Polysciences #86057) at RT for 10mins. Nuclei were blocked for 5 min at RT then incubated in 1:100 dilution of primary antibody for 2 hr at 4°C, washed, incubated in a 1:200 dilution of pAG MNase for 1 hr at 4°C, and washed again. The bound MNase was activated with 2 mM CaCl_2_ and allowed to digest for 30 mins, then stopped using 2x STOP buffer (200 mM NaCl, 20 mM EDTA, 4 mM EGTA, 50 μg/mL RNase A, 40 μg/mL glycogen). Nuclei were incubated at 37°C for 20 min followed by centrifuging for 5 min at 16,000xg, drawing off the DNA fragments with the supernatant. The extracted fragments were treated with SDS and proteinase K at 70°C for 10 min followed by phenol chloroform extraction. Purified DNA was resuspended in 50 μL of water and verified by Qubit dsDNA high sensitivity and Fragment Analyzer. Antibodies used were: H3K4me3, Invitrogen #711958, Lot #2253580; H3K27ac, ActiveMotif #39135, Lot #06419002; V5, Invitrogen #R960-25, Lot #2148086.

For transcription factor CUT&RUN, *pou5f3.3.L* and *sox3.S* IVT templates were cloned from cDNA using primers for *pou5f3.3.L* – NM_001088114.1 (F: GGACAGCACGGGAGGCGGGGGATCCGACCAGCCCATATTGTACAGCCAAAC; R: TATCATGTCTGGATCTACGTCTAGATCAGCCGGTCAGGACCCC) and *sox3.S* - NM_001090679.1 (F: TATAGCATGTTGGACACCGACATCA; R: TTATATGTGAGTGAGCGGTACCGTG) into N-terminal V5-pBS entry plasmids using HiFi assembly (NEB #E2621) for *pou5f3.3* and BamHI/XbaI for *sox3*. IVT was done using NEB HiScribe T7 ARCA kit (#E2065S) on NotI-linearized plasmid for 2hrs at 37°C, then treated with 5U of TURBO DNaseI (Invitrogen #AM2238) for 15 min. mRNA was purified using NEB Monarch RNA Cleanup Columns (#T2030) and stored at −80°C until use. For injection, immediately after dejellying, stage 1 embryos were placed in 4% Ficoll-400 in MR/3. Each embryo was injected with 5 nL of 40 ng/μL of mRNA on opposite sides, for a total of 10 nL per embryo. Factor-specific no-antibody CUT&RUN samples were made using the same injected embryos.

CUT&RUN libraries were constructed using the NEB Ultra II DNA library prep kit (NEB #E7645) according to manufacturer’s protocol. DNA was end repaired and then ligated to sequencing adaptors diluted 1:10. Ligated DNA was purified with 0.9x AmpureXP beads and PCR amplified for 15 cycles, then purified again with 0.9x AmpureXP beads. Libraries were size selected to 175 – 650 bp via 1.5% TBE agarose gel and gel purified using the NEB Monarch DNA gel extraction kit (#T1020) before being verified by Qubit dsDNA high sensitivity and Fragment Analyzer prior to multiplexed paired-end sequencing on an Illumina NextSeq 500 at the Health Sciences Sequencing Core at Children’s Hospital of Pittsburgh.

### ATAC-seq

ATAC procedure was from Esmaeili et al (Esmaeili et al., 2020) Embryos were grown in MR/3 until desired NF stage and devitellinized individually with fine watch-maker forceps. Ectodermal explants (animal caps) were dissected using watch-maker forceps in 0.7x MR. Two caps were transferred to 1 mL of ice-cold PBS and centrifuged at 500xg in 4°C for 5 min twice. After washing with PBS, caps were lysed in 50 μl of RSB buffer (10 mM Tris pH 7.4, 10 mM NaCl, 3 mM MgCl2, 0.1% Igepal CA-630) with a clipped P200 pipet. The lysate was centrifuged again for 10 min and the supernatant was drawn off. The pellet was resuspended in 47.5 μl TD buffer (10mM Tris pH 7.6, 5 mM MgCl2, 10% dimethylformamide) and 2.5 μl of 3 μM transposome (see below) was added. Nuclei were transposed with gentle shaking for 1 hr at 37° C before adding 2.5 μl proteinase K and incubating overnight at 37°C. Transposed DNA was purified using EconoSpin Micro columns (Epoch) and amplified using 25 μM indexed Nextera primers with Thermo Phusion Flash master mix for 12 cycles. Primers used were: CAAGCAGAAGACGGCATACGAGAT[i7]GTCTCGTGGGCTCGG with i7 indices 707 – gtagagag; 714 –tcatgagc; 716 – tagcgagt; and AATGATACGGCGACCACCGAGATCTACAC[i5]TCGTCGGCAGCGTC with i5 indices 505 – gtaaggag; 510 – cgtctaat; 517 – gcgtaaga; 520 – aaggctat. The amplified library was column cleaned and verified by Qubit dsDNA high sensitivity and Fragment Analyzer and sequenced multiplexed paired end at the Health Sciences Sequencing Core at Children’s Hospital of Pittsburgh. After initial sequencing, libraries were subsequently size selected on an agarose gel to enrich for 150-250 and 250-600 bp fragments and resequenced pooled.

Transposomes were constructed according to Picelli et. al. (Picelli et al., 2014) Adapter duplexes for Tn5ME-A (TCGTCGGCAGCGTCAGATGTGTATAAGAGACAG) + Tn5MErev ([phos]CTGTCTCTTATACACATCT) and Tn5ME-B (GTCTCGTGGGCTCGGAGATGTGTATAAGAGACAG) + Tn5MErev were each annealed in 2 μl of 10X annealing buffer (100 mM HEPES pH 7.2, 500 mM NaCl, 10 mM EDTA) using 9 μl of each oligo at 100 μM, heated to 95°C for 1 min then ramped down to 25°C at 0.1°C/s in a thermocycler. The two duplexes were held at 25°C for 5 min then mixed together. On ice, 35 μl of hot glycerol was cooled to 4°C then 35 μl of the primer mixture and 25 μl of Tn5 (Addgene #112112) was added and mixed and held at 1 hr at RT with gentle pipet mixing every 15 min. Transposomes were stored at −20°C.

### Transcriptomic analysis

RNA-seq reads were mapped to the *X. laevis* v9.2 genome using HISAT2 v2.0.5 (Kim et al., 2015) (--no-mixed --no-discordant). Mapped reads were assigned to gene exons (Xenbase v9.2 models) using featureCounts v2.0.1 (in reversely-stranded paired-end mode with default parameters, and to introns with --minOverlap 10 on a custom intron annotation: starting with all introns from the v9.2 GFF file, subtract (a) all regions detected in stage 5 RNA-seq at >2 read coverage, strand specifically; (b) all regions that overlap an annotated exon from a different transcript form; (c) regions that overlap repetitive elements as defined by RepeatMasker (UCSC) and Xenbase-annotated transposons, not strand specifically; (d) regions that ambiguously map to more than one distinct gene’s intron (i.e., transcript forms of the same gene are allowed to share an intron, but not between different genes).

DESeq2 v4.0.3 (Love et al., 2014) was used for statistical differential expression analysis. To build the DESeq2 model, exon and intron raw read counts were treated as separate rows per gene in the same counts matrix (intron gene IDs were preceded with a “i_” prefix). Only genes annotated by Xenbase as “protein_coding,”“lncRNA,” or “pseudogene” were retained. Low-expressed genes were removed (exon reads per million (RPM) < 0.5 across all samples) and then low-depth intron features were removed (intron raw read count ≤ 10 or reads per kilobase per million (RPKM) < 0.25 across all samples). Comparisons were made between batch-matched samples where possible, to account for variations in the maternal contribution between mothers. Significant differences with adjusted p < 0.05 and log2 difference ≥ 1.5 were used for downstream analysis. High-confidence activated genes had significant increases in DMSO vs Triptolide for both batches and stage 9 vs stage 5. High-confidence primary-activation “first-wave” genes were high-confidence activated and had significant increase in DMSO vs Cycloheximide. Homeologous genes were paired according to Xenbase GENEPAGE annotations. Genes were considered maternal if they had average stage 5 TPM ≥ 1. To calculate magnitude of effect for graphing and sorting, the maximal ļlog2 fold differenceļ of average exon TPM and average intron RPKM was chosen per gene.

For mir-427 gene identification and RNA-seq coverage visualization, miRBase (Kozomara et al., 2019) hairpin sequences MI0001449 and MI0038331 were aligned to the v9.2 and v10.1 reference genomes using UCSC BLAT (Kent, 2002) and maximal possible read coverage was graphed allowing all multimappers. To align the v10.1 Chr1L and Chr1S regions flanking the Chr1L mir-427 locus, genomic sequence was extracted between homeologous genes upstream and downstream mir-427. Local alignments with E-value < 1e-10 were retained from an NCBI BLAST 2.11.0+ blastn alignment (Camacho et al., 2009).

dN/dS ratios were calculating using PAML v4.9f (Yang, 1997) with L-S pairwise CDS alignments produced by pal2nal v14 (Suyama et al., 2006) on amino-acid alignments by EMBOSS needle v6.6.0.0 (-gapopen 10 -gapextend 0.5) (Rice et al., 2000).

All other statistical tests were performed using R v4.0.4 (R. Core Team, 2013).

### Chromatin profiling analysis

CUT&RUN and ATAC-seq paired-end reads were mapped to the *X. laevis* v9.2 genome using bowtie2 v2.4.2 (Langmead and Salzberg, 2012) (--no-mixed --no-discordant) and only high-quality alignments (MAPQ ≥ 30) were retained for subsequent analysis. Read pairs were joined into contiguous fragments for coverage analyses. For transcription factor CUT&RUN, reads were trimmed using trim_galore v0.6.6 and Cutadapt v1.15 (Martin, 2011) in paired-end mode (--illumina --trim-n). Downstream analyses were performed using custom scripts with the aid of BEDtools v2.30.0 (Quinlan and Hall, 2010), Samtools v1.12 (Li et al., 2009), and deepTools v3.5.1 (Ramírez et al., 2014).

For promoter-centered analyses, one transcript isoform per gene was selected from Xenbase 9.2 annotations: the most upstream TSS with non-zero RNA-seq coverage at Stage 9 was used, otherwise the most upstream TSS if no RNA-seq evidence.

To identify open chromatin regions, aligned ATAC-seq fragments pooled between replicates were filtered to <130 bp, then peaks called using MACS2 v2.2.7.1 (Zhang et al., 2008) with an effective genome size of 2.4e9 (number of non-N bases in the reference sequence). CUT&RUN no-antibody samples were used as the control sample. To further exclude probable falsepositive regions, peaks overlapping any of the following repetitive regions were removed: (a) scRNA, snRNA, snoRNA, or tRNA as annotated by Xenbase; (b) rRNA as determined by RepeatMasker and BLASTed 45S, 16S, 12S, and 5S sequences. Peaks on unassembled scaffolds were also excluded.

Putative enhancers had 2-fold enriched pooled stage 8 H3K27ac CUT&RUN coverage over no antibody, ≥1 RPM H3K27ac coverage, and <0.5 RPM no antibody coverage, in a 500-bp window centered on ATAC-seq peak summits. Enhancers were classified as distal if they were > 1 kb from any Xenbase 9.2 annotated TSS, proximal otherwise.

For transcription factor peak calling, the Sox3 sample was down-sampled to match Pou5f3 read depth (~12 M read pairs) using samtools view -s. No-antibody samples were pooled as a uniform background. MACS2 was run as above, and SEACR v1.3 (Meers et al., 2019) was run in norm relaxed mode. Peak calls were not used for enhancer analyses; rather, enhancers or homeologous regions with ≥1 RPM CUT&RUN coverage and ≥2-fold enrichment over no antibody in a 200-bp window were considered bound.

Coverage heatmaps were generated using deepTools on reads-per-million normalized bigWigs or enrichment over no-antibody bigWigs generated using deepTools bigwigCompare (-operation ratio --pseudocount 0.1 --binSize 50 --skipZeroOverZero).

For density heatmaps, enhancer pairs were annotated as differential or conserved based on one or both partners, respectively, mapping to a putative enhancer, as described above. The total region of each putative enhancer corresponding to ≥2-fold H3K27ac enrichment was calculated and converted to a bigWigs representing the genomic location of each enriched region. Pairs were similarly annotated as differentially or both TF bound based on ≥2-fold enrichment over no antibody for either TF at one or both partners, respectively, and converted to bigWigs representing the genomic location of each bound putative enhancer. Density heatmaps were generated as above and plotted with respect to selected TSSs.

### Motif finding

Enriched sequence motifs in enhancers were identified using Homer v4.11.1 (Heinz et al., 2010) in scanning mode against the vertebrate database, using 200 bp of sequence centered on the ATAC-seq peak for enhancers and 500 bp of sequences centered on the TSS for promoters. Enrichment was calculated using one set of homeologous regions (L or S) as the foreground and the other as the background. The top representative motif per DNA binding domain was reported. For transcription factor peaks, Homer was used in de novo mode on the top 500 peaks. The top motif was extracted for each of Pou5f3 and Sox3, then scanned against the entire set of peaks.

### Homeologous enhancer identification

Each chromosome pair (e.g., chr1L and chr1S) was aligned using lastZ-1.04.00 (Harris, 2007) and UCSC Genome Browser utilities (Kent et al., 2002) with parameters adapted from the UCSC Genome Browser previously used to align *X. tropicalis* with *X. laevis* (http://www.bx.psu.edu/miller_lab/dist/README.lastz-1.02.00/README.lastz-1.02.00a.html; http://genomewiki.ucsc.edu/index.php/XenTro9_11-way_conservation_lastz_parameters) (no automatic chaining; open=400, extend=30, masking=0, seed=1 {12of19}, hspthreshold=3000, chain=0, ydropoff=9400, gappedthreshold=3000, inner=2000). Chaining and netting were done with axtChain linearGap set to medium and chainSplit lump=50. Nets were generated using default chainNet and the highest scoring chains were selected from those nets using default netChainSubset. Reciprocal best chains were identified according to UCSC Genome Browser guidelines. The highest scoring chains were reverse referenced, sorted, and then converted to nets using default chainPreNet and chainNet (-minSpace=1 -minScore=0). Reciprocal best nets were selected with default netSyntenic. The new highest scoring best chains were extracted using netChainSubset, converted back to the original reference, and netted as described prior, resulting in reciprocal best, highest scoring chains for use with liftOver.

In the first pass, 500-bp enhancer regions centered on the ATAC-seq peak were lifted to the homeologous subgenome with a 10% minimum sequence match requirement. For enhancers that failed this liftOver, 5-kb enhancer regions were lifted over; as a stringency check, each 2.5-kb half was also individually lifted over, and only regions correctly flanked by both halves were retained. If an enhancer’s homeologous region also overlaps an annotated enhancer, it was considered conserved, otherwise it was considered subgenome-specific. To test synteny, the 5 closest Xenbase-annotated genes up- and downstream of each region in a homeologous pair were compared.

### Comparison with *X. tropicalis* and zebrafish

*X. tropicalis* wild-type RNA-seq reads from Owens et al (Owens et al., 2016), RiboZero stage 5 (SRA: SRR1795666) and stage 9 (SRA: SRR1795634), were aligned by HISAT2 as above and mapped to Xenbase v10 gene annotations using featureCounts. Pou5f3/Sox3 morpholino and alpha-amanitin-affected genes were obtained from published data tables from Gentsch et al (Gentsch et al., 2019), and the JGI gene accession numbers were mapped to Xenbase GenePage IDs (v7.1). Significantly affected genes were 1.5-fold decreased and adjusted p < 0.05. Genes with TPM > 1 at either stage 5 or stage 9 were considered embryonic expressed.

Zebrafish annotations for activated and Pou5f3 / Nanog / SoxB1 affected genes were obtained from Lee & Bonneau et al (Lee et al., 2013) and associated to Xenopus genes using Ensembl ortholog annotations (Xenbase to Zfin). First-wave activated zebrafish genes are significantly increased in the U1/U2 spliceosomal RNA inhibited sample over alpha-amanitin (DESeq2 adjusted p < 0.05), activated genes are significantly increased by 6 h.p.f. over alpha-amanitin. Pou5f3/SoxB1 affected genes were significantly decreased in the Pou5f3-SoxB1 double loss of function versus wild-type. Nanog-affected genes were significantly decreased in triple loss of function (NSP) but not Pou5f3-SoxB1 double loss of function. Genes with TPM > 1 at 2, 4, or 6 h.p.f. were considered embryonic expressed.

To identify putative conserved enhancers in *X. tropicalis, X. laevis* enhancers were lifted over as above to the *X. tropicalis* v9.2 genome using liftOver chains from the UCSC Genome Browser (xenLae2ToXenTro9, 10% minimum sequence match). Successfully lifted over regions were intersected with published *X. tropicalis* H3K27ac stage 9 peaks from Gupta et al (Gupta et al., 2014) that were lifted from the *X. tropicalis* v2 genome to the v9 genome, passing through v7 and requiring 90% minimum sequence match, using liftOver chains from UCSC Genome Browser (xenTro2ToXenTro7 and xenTro7ToXenTro9). *X. laevis* enhancers were lifted over to the zebrafish GRCz11 genome using liftOver chains from the UCSC Genome Browser, passing through *X. tropicalis* (xenLae2ToXenTro9, 10% minimum sequence match; then xenTro9ToXenTro7, 90% minimum sequence match, then xenTro7ToDanRer10, 10% minimum sequence match, then danRer10ToDanRer11 requiring 90% minimum sequence match). Acetylation at zebrafish dome stage was then assessed by intersecting with H3K27ac ChIP-seq peaks from Bogdanovic et al (Bogdanovic et al., 2012) (GEO: GSM915197): reads were aligned to the GRCz11 genome using bowtie2 as above, and peaks called using macs2 as above with an effective genome size of 4.59e8 and no control sample.

