## Supplementary Figures for "Hybridization led to a rewired pluripotency network in the allotetraploid *Xenopus laevis*"

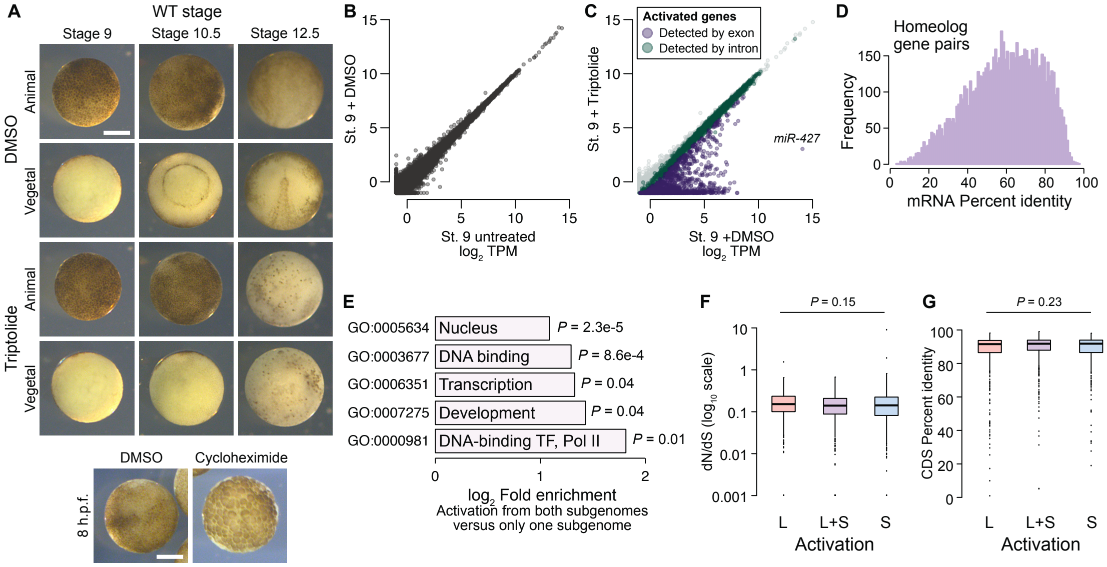


**Supplementary Fig 1. Measuring genome activation.** (**A**) (Top) Animal and vegetal views of embryos treated with DMSO (vehicle) versus triptolide. Triptolide-treated embryos fail to gastrulate. (Bottom) Comparison of DMSO versus cycloheximide treated embryo. Treatment was at stage 8, which inhibits progression to stage 9. Scale bar = 0.5 mm. (**B**) Biplot of RNA-seq for untreated versus DMSO-treated embryos at stage 9, showing no effect on the transcriptome. (**C**) Biplot of RNA-seq for DMSO versus triptolide treated embryos, showing inhibited activation as detected by exonic (purple) and intronic (green) signal. The predicted *mir-427* primary transcript is labeled and exhibits >95% expression inhibition. (**D**) Histogram of mRNA percent identity between homeolog pairs, as measured by Needleman-Wunsch alignment. Maximum is 0.975 (5 in every 200 bases, which should be generally distinguishable by RNA-seq using 2x100 sequencing reads) (**E**) Significantly (FDR < 0.05, Fisher's exact test, two-sided) enriched Gene Ontology terms in genes activated from both homeologs, as compared to genes activated from only one subgenome. (**F**) Boxplots of non-synonymous to synonymous substitution rate ratio (dN/dS) shown on a log10 scale, for genes activated from both subgenomes or only one subgenome (*P* = 0.15, Kruskal-Wallis test; median L = 0.15, LS = 0.14, S = 0.14). All gene groups trend toward stabilizing selection. (**G**) Boxplots of CDS percent similarity for activation groups (*P* = 0.23, Kruskal-Wallis test). For all boxplots: center line, median; box limits, upper and lower quartiles; whiskers, 1.5x interquartile range; points, outliers.

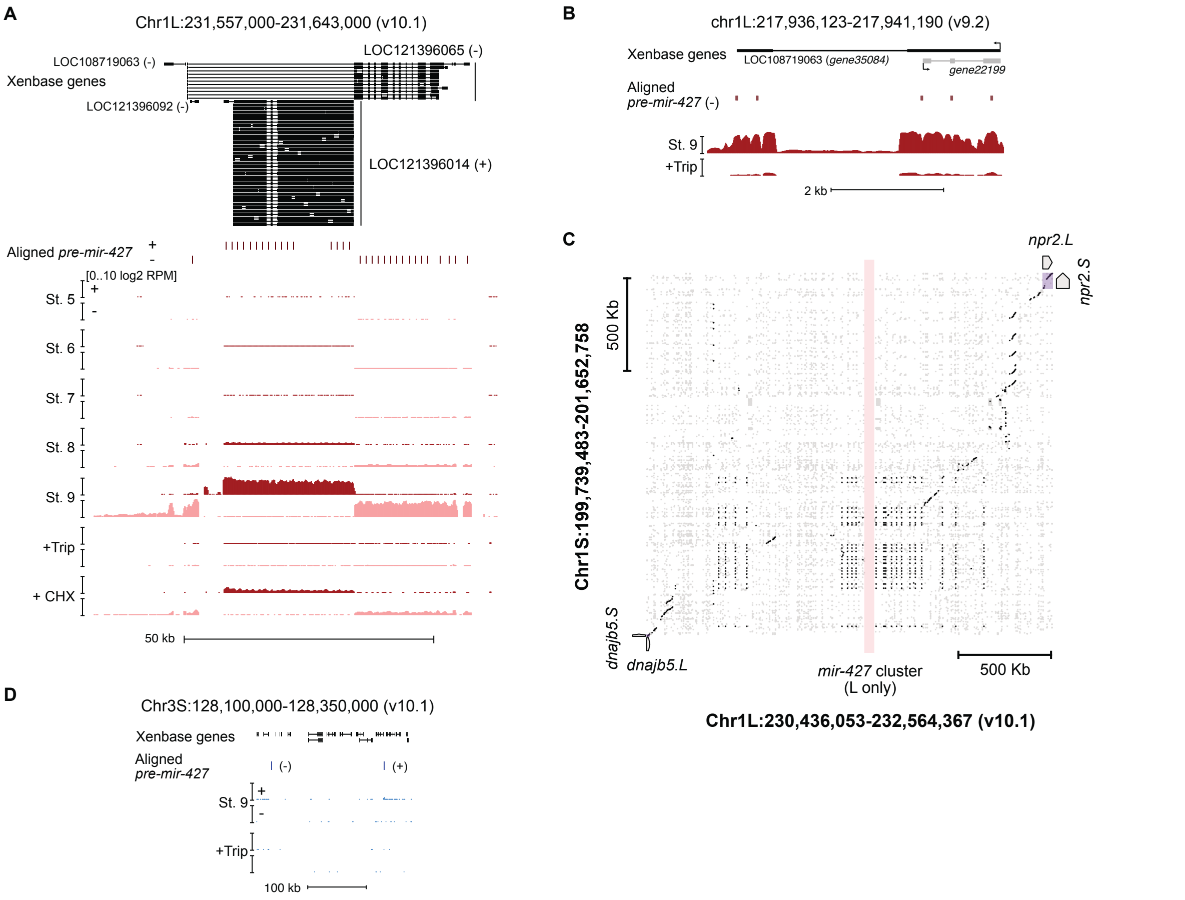


**Supplementary Fig 2. The mir-427 locus.** (**A**) Browser tracks showing strand-separated log2 reads-per-million RNA-seq coverage over the predicted *mir-427* primary transcript near the telomere of Chr1L on the v10.1 genome assembly. Xenbase gene isoforms are annotated at the top, aligned precursor mir-427 sequences are annotated in the middle according to strand orientation. (**B**) Browser track showing log2 reads-per-million RNA-seq coverage over the presumed mir-427 encoding region on the v9.2 genome assembly. The overlapping antisense transcript is not transcribed (all coverage shown is sense to the *mir-427* transcript). (**C**) Dot matrix alignment plot showing BLAST local alignments between v10.1 Chr1L and Chr1S in the region flanking the mir-427 locus. Repetitive sequence alignments (Xenbase soft-masked genomic sequence) are shown in gray, non-repetitive alignments in black. Upstream (*dnajb5*) and downstream (*npr2*) homeologous genes are labeled. The L-specific mir-427 locus is highlighted in light red, showing no alignments to Chr1S. (**D**) Region of v10.1 Chr3S where two additional sequence matches to the mir-427 hairpin are found by BLAT. However, there is minimal RNA-seq coverage, suggesting the Chr1L locus is the only bona fide *mir-427* encoding region in the v10.1 assembly. Log2 reads-per-million coverage is shown on the same scale as panel (A).

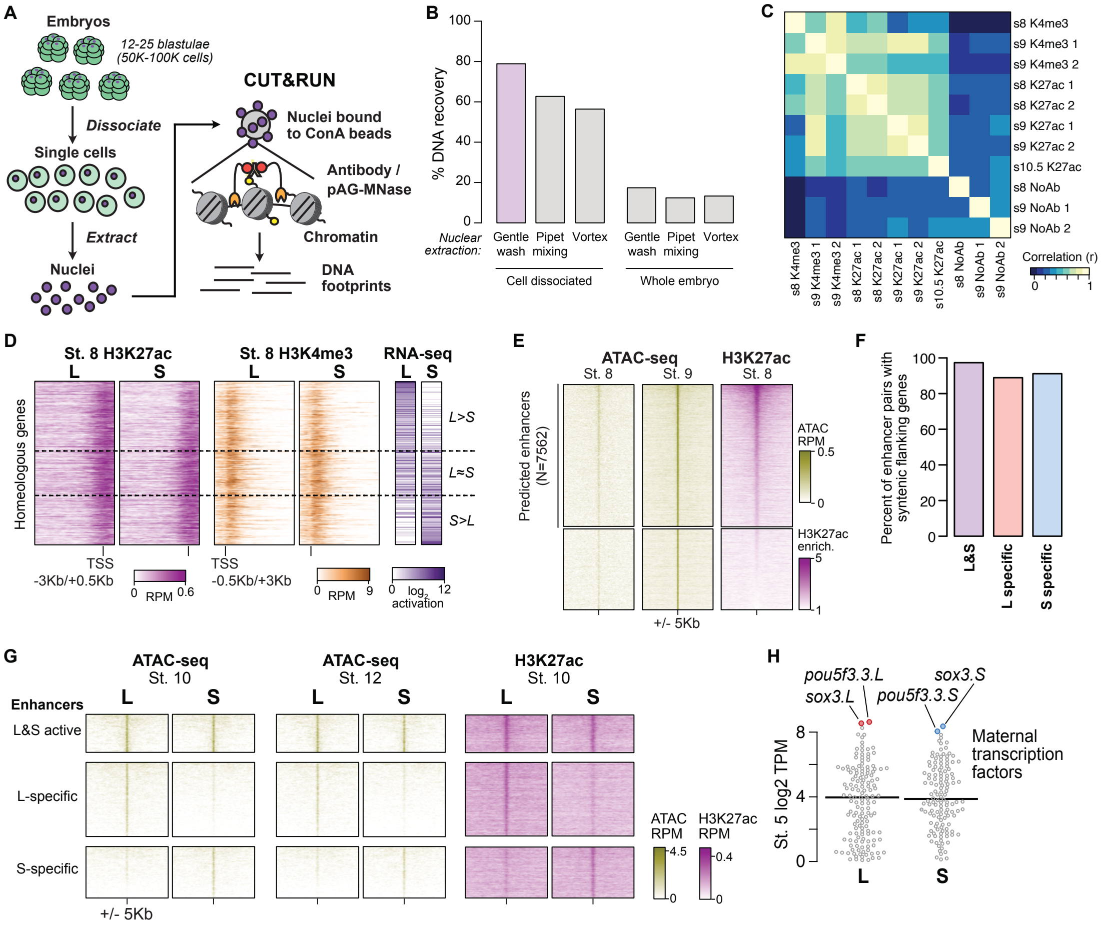


**Supplementary Fig 3. Profiling homeologous regulatory elements.** (**A**) CUT&RUN for *X. laevis* blastulae requires cell dissociation prior to nuclear extraction. (**B**) Comparison of different nuclear extraction techniques. Percent DNA recovered was estimated by NanoDrop quantification of phenol-chloroform extracted DNA after nuclear extraction, as a percentage of theoretical total nuclear DNA mass based on the length of the reference genome sequence. Three nuclear extraction methods were tested with and without cell dissociation: gentle washing by pipeting buffer on the surface of the cells, pipet mixing, vortexing at 1500 rpm. (**C**) Heatmap of pairwise sample correlation between CUT&RUN samples, as measured by log2 coverage in a 1 Kb window around the center of ATAC-seq open regions (N = 41083). (**D**) CUT&RUN coverage over paired homeologous gene regions around the TSS. Gene pairs are sorted according to L versus S RNA-seq activation ratio (right). (**E**) Heatmaps showing ATAC-seq peaks divided into putative enhancers with H3K27ac CUT&RUN enrichment, versus non-enhancers lacking H3K27ac. (**F**) Proportion of predicted homeologous enhancers that are flanked upstream and downstream by homeologous genes (at least 1 of the 5 nearest genes up/downstream). (**G**) Heatmaps of later-stage ATAC-seq (Esmaeili et al 2020) and H3K27ac CUT&RUN coverage (this study) plotted over homeologous enhancers. (**H**) Maternal (stage 5) RNA-seq levels for L and S sequence-specific transcription factors, as annotated by Gene Ontology. *pou5f3.3* and *sox3* are the top expressed transcription factors for both L and S.

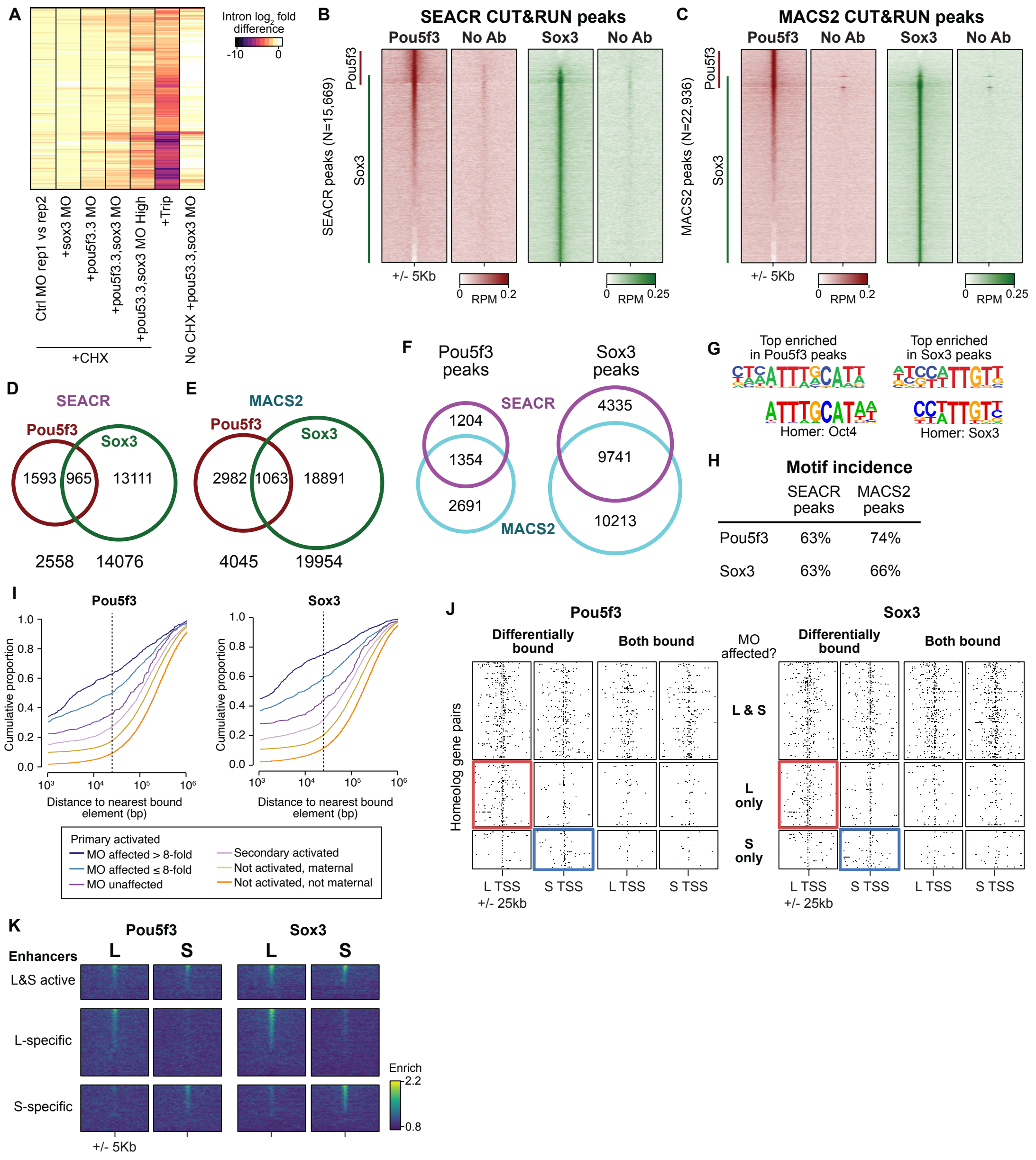


**Supplementary Fig 4. Assessing Pou5f3 and Sox3 roles in genome activation.** (**A**) Heatmap showing RNA-seq intron log2 fold difference compared to control for intron-containing activated genes. Column 1 compares replicates of cycloheximide (CHX) treated embryos injected with GFP control morpholino (MO); columns 2-5 show cycloheximide-treated embryos injected with individual sox3 and pou5f3.3 morpholino and both sox3 + pou5f3.3 morpholino at lower and higher concentrations compared to their respective control GFP morpholino injected embryos; column 6 shows triptolide-treated embryos compared to DMSO control; and column 7 shows embryos treated with sox3 + pou5f3.3 morpholino without cycloheximide compared to GFP morpholino. All samples are from embryos collected when untreated wild-type controls were at stage 9. (**B, C**) Heatmaps of Pou5f3.3 and Sox3 CUT&RUN coverage over SEACR (C)

and MACS2 (D) predicted peaks. The union of peaks per method is shown. Pou5f3 and Sox3 signal overlaps across most of the peaks, though it appears that both methods under-call Pou5f3 peaks. No Ab = no antibody. (**D, E**) Venn diagrams showing peak overlap between factors for SEACR (E) and MACS2 (F). (**F**) Venn diagrams showing peak overlap between methods. (**G**) Top enriched motif for Homer de novo motif finding for each factor compared to the closest database match. (**H**) Table of motif occurrence of the de novo identified Pou5f3 and Sox3 motifs among the peaks. (**I**) Cumulative distributions of distance from a Pou5f3-bound (left) or Sox3-bound (right) regulatory element. Curves represent gene groups according to the degree that they are affected by pou5f3.3/sox3 morpholino treatment. (**J**) Maps showing density of Pou5f3-bound (left) and Sox3-bound (right) regulatory elements around paired homeologous TSSs, divided into elements with differential homeologous L&S binding (i.e., one bound, the other not) versus both L & S homeologous region bound. TSSs are grouped according to whether both L & S homeologs are affected by pou5f3.3/sox3 morpholino treatment, or only one or the other homeolog is affected. For Pou5f3, $P = 1.5\times10^{-5}$, Kruskal-Wallis test for L-S differential bound enhancer count difference among the three gene groups; for Sox3, $P = 1.8\times10^{-5}$. For both-bound enhancers, $P = 0.13$ for Pou5f3, $P = 0.067$ for Sox3. (**K**) Heatmaps showing Pou5f3 and Sox3 CUT&RUN binding enrichment over no-antibody control, plotted over homeologous enhancers as previously defined by ATAC-seq and H3K27ac.

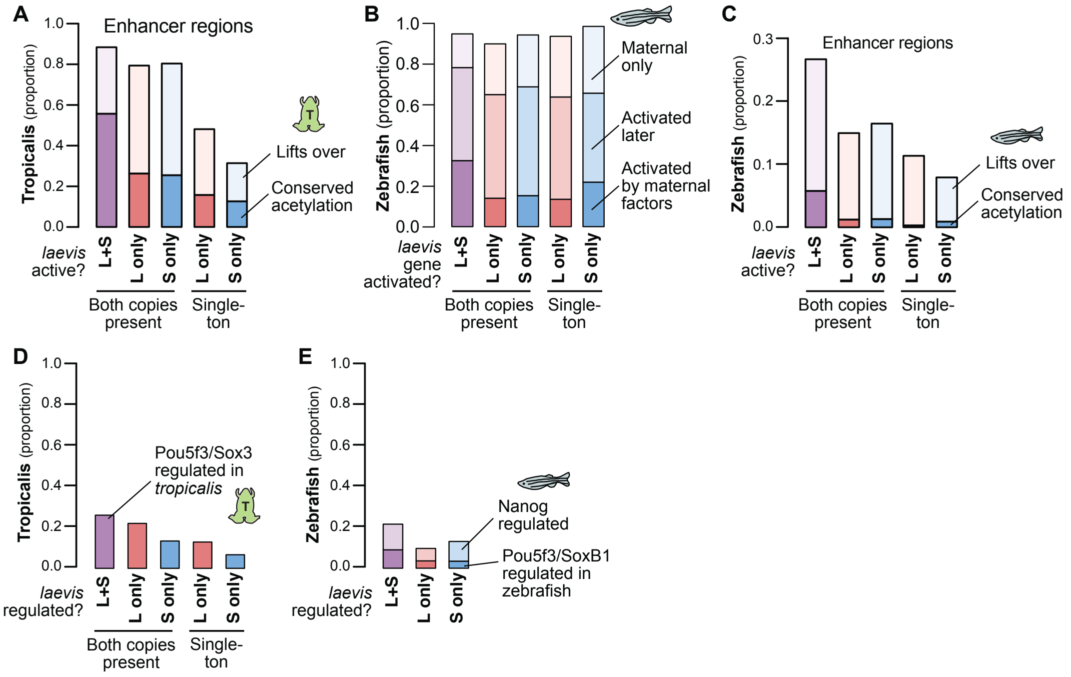


**Supplementary Fig 5. Shared patterns of activation with other taxa.** (**A**) Similar to Fig. 4C: barplots showing the proportion of *X. laevis* enhancers across activity categories that are acetylated in *X. tropicalis*, additionally showing the proportion of enhancers that lift over but are not acetylated in *X. tropicalis*. (**B**) Barplots showing the proportion of *X. laevis* genes in different homeolog activation categories whose orthologs are also activated in zebrafish as part of the first wave by maternal factors, activated by 6 h.p.f., or part of the maternal contribution. Both-activated homeologs are more likely to also be activated in zebrafish in the first wave ($P = 8.0\text{x}10^{-12}$, $\chi$-squared test, 4 d.o.f.). (**C**) Barplots showing the proportion of enhancers that lift over and are acetylated in zebrafish according to Bogdanovich et al. 2012. L+S conserved enhancers have low conservation with zebrafish, but significantly higher proportion than L- or S-only enhancers ($P = 9.4\text{x}10^{-80}$, $\chi$-squared test, 4 d.o.f). (**D**) Barplots showing the proportion of Pou5f3/Sox3-regulated *X. laevis* genes also regulated by Pou5f3/Sox3 in *X. tropicalis* according to Gentsch et al. Both-regulated homeologs are more likely to also be regulated in *X. tropicalis* ($P = 4.2\text{x}10^{-4}$, $\chi$-squared test, 4 d.o.f.). (**E**) Barplots showing the proportion of Pou5f3/Sox3-regulated *X. laevis* genes also regulated by Nanog/Pou5f3/Sox3 in zebrafish according to Lee & Bonneau et al. 2013. Both-regulated homeologs are more likely to also be regulated by Pou5f3/SoxB1 in zebrafish ($P = 0.0096$, $\chi$-squared test, 2 d.o.f.), but also more likely to be regulated by Nanog in zebrafish ($P = 3.4\text{x}10^{-4}$, $\chi$-squared test, 2 d.o.f.).

**Supplementary Table 1**
RNA-seq expression values and DESeq2 comparisons

**Supplementary Table 2**
Annotations and expression values for activated genes

**Supplementary Table 3**
Transcription start site coordinates used for all genes

**Supplementary Table 4**
Motif search results

**Supplementary Table 5**
Enhancer annotations

**Supplementary Table 6**
Comparative transcriptomics with *X. tropicalis* and zebrafish
